# Transcriptomic dissection of Intraepithelial Papillary Mucinous Neoplasms progression by spatial technologies identified novel markers of pancreatic carcinogenesis

**DOI:** 10.1101/2022.10.12.511894

**Authors:** Antonio Agostini, Geny Piro, Frediano Inzani, Giuseppe Quero, Annachiara Esposito, Alessia Caggiano, Lorenzo Priori, Alberto Larghi, Sergio Alfieri, Raffaella Casolino, Vincenzo Corbo, Andrew V Biankin, Giampaolo Tortora, Carmine Carbone

## Abstract

Intraductal papillary mucinous neoplasms (IPMN) are one of the main precursor lesions of Pancreatic Ductal Adenocarcinoma (PDAC). The number of patients diagnosed with IPMN is constantly increasing. While in most of the cases IPMN present as indolent and nonmalignant entities, some degenerate into PDAC. The main mechanisms behind the IPMN progression to malignancy is still not fully understood.

This is mainly due to the technological limit of the analyzes and to cysts heterogeneity whose malignant transformation potential is estimated based on size and degree of dysplasia without take in consideration the transformation time and therefore the real malignancy potential.

Moreover, there is a general lack of consensus diagnostic markers to discern the Low-grade nonmalignant from High-grade malignant IPMN. In this study, we used two different Spatial Transcriptomic technologies (Visium, and GeoMx) to investigate the transcriptome of Low-grade dysplasia nonmalignant IPMN, Borderline IPMN, and High-grade dysplasia malignant IPMN to dissect the main mechanism that drives carcingenesis and to find specific markers associated to risk of tumor progression.

We performed Visium spatial transcriptomics on two TMAs containing three Low-grade dysplasia nonmalignant IPMN, one Borderline IPMN, two High-grade dysplasia malignant IPMN, and two PDAC.

We identified three specific epithelial cell clusters that characterize Low-grade dysplasia IPMN, Borderline IPMN, and High-grade dysplasia malignant IPMN and three transcription factors whose expression is associated with each grade. High-grade malignant IPMN were characterized by high expression levels of *NKX6-2* and other markers of gastric isthmus cell lineage such as *MUC5AC, PSCA, FERIL6.* The *SPDEF* high IPMN cluster was found in Borderline IPMN and spotted in some regions of High-grade malignant IPMN. This cluster was characterized by high expression levels of *SPDEF* and other goblet cell lineage markers such as *TFF2, AQP5,* and *MUC6.* Low-grade nonmalignant IPMN were characterized by high expression levels of *HOXB3, HOXB5, ZNF117.* The association of these markers with the different grades was validated by GeoMx spatial transcriptomics on 43 additional IPMN samples divided according to their grade of dysplasia and malignancy.

To better understand the transcriptomic changes along IPMN progression we performed spatial trajectory inference and we found that *SPDEF* high IPMN cluster cells are likely to evolve into *NKX6-2* high malignant IPMN, and we found that this switch is characterized by the expression of *NKX6-2* and other gastric markers.

Taken together, the results presented here not only shed more light in to IPMN and PDAC oncogenesis, but also provided a plethora of novel malignancy-associated markers to be tested in diagnostic routine, to better delineate IPMN progression in patients and improve clinical management.

## Introduction

Intraductal papillary mucinous neoplasms (IPMN) are cystic dilatation of the ductal system with papillary projections characterized by a mucin-producing epithelium and classified as premalignant lesions (1,2). The number of patients diagnosed with IPMN is constantly increasing, reaching around 8% of the world population (2,3). The IPMNs have different neoplastic transformation potential based on the specific dysplasia grade (high-grade dysplasia (HGD) or low-grade dysplasia (LGD)), on the types of the origin of the epithelial precursors (4) and on the localization site (main duct, MD, or branch ducts, BD). Surgery is recommended for IPMN patients with HGD or for cysts in the MD of the pancreas (5). For the other part of the patients a follow-up is essential. However, IPMN surgical treatment is invasive and sometimes followed by complications (6).

In a recent retrospective multicenter study of IPMN patients subjected to surgery (n = 1074) the 14.4% developed postoperative recurrence, and that 34.4% of patients operated on for high-risk IPMN developed lesions over 5 years after surgery (7). Of the Low-risk IPMN patients who were assigned to clinical follow up, around 1-11% developed pancreatic ductal adenocarcinoma (PDAC) (8,9).

Since the pancreas is a highly deadly disease (5 years OS < 8%) with a high recurrence rate and projected to be the second cancer-related death in US (10) it is a urgent need to define the molecular markers capable of predicting which cysts need surgery and to detect the early mechanisms responsible for their neoplastic transformation, both for prevention and for therapeutic intervention.

Many papers tried to define the biology and prognosis of invasive IPMNs, using data based on histological and precursor epithelial subtypes (11,12) and on genetic alterations, mainly KRAS, GNAS, and RNF43 or mutations such as TP53, CDKN2A, SMAD4 that do not associate however to the degree of dysplasia nor to the histological subtype. Studies based on IPMN tissue bulk transcriptomic and proteomic analysis of the fluid cysts attempted to identify transformation markers capable of predicting the malignant transformation potential, however none of the works to date have produced results that have actually been translated into clinical use.

This is mainly due to the technological limit of the analyzes and to cysts heterogeneity whose malignant transformation potential was based exclusively on the size and dysplasia degree and in which the transformation time and therefore the real malignancy potential were not taken into consideration.

In this study, using two spatial transcriptomic technologies we identified and validated 3 gene clusters with prognostic value. These clusters could discriminate between IPMNs with different neoplastic transformation potential distinguishing low-risk IPMN from high-risk IPMN and will therefore allow the development of a molecular risk stratification test for patients with IPMN. Furthermore, here we unveil the factors activated along the neoplastic transformation of IPMN by tracing the route that leads from the IPMN to the invasive cancer.

## Methods

### Patient Material

We gathered six Low-grade (LG) and one Borderline (Br) IPMNs from patients with a follow up of more than 10 years that never developed PDAC. From these series we chose for Visium Spatial transcriptomics (10X Genomics, USA) three LG IPMN, one Br IPMN, and we analyzed them together with two High-grade (HG) IPMN, and 2 PDAC. All FFPE samples were selected after proper evaluation from an expert pathology of dysplasia, progression stage, and cellularity (>30%). All these samples had a good RNA quality with a DV200 score (> 50%). Two Tissue Macro Array (TMA) (Fig.1) were build using 1,5 core punches and the robotic TMA builder Galileo (ITS, Italy): TMA1 containing two LG IPMN, one HG IPMN and one PDAC; and TMA2 containing one LG IPMN, one Borderline IPMN, one HG IPMN and one PDAC (Fig 1). TMAs were conserved at 4°C enclosed in a sealed bag with silica gel beads to preserve RNA integrity and avoid oxidation.

**Figure 1.**
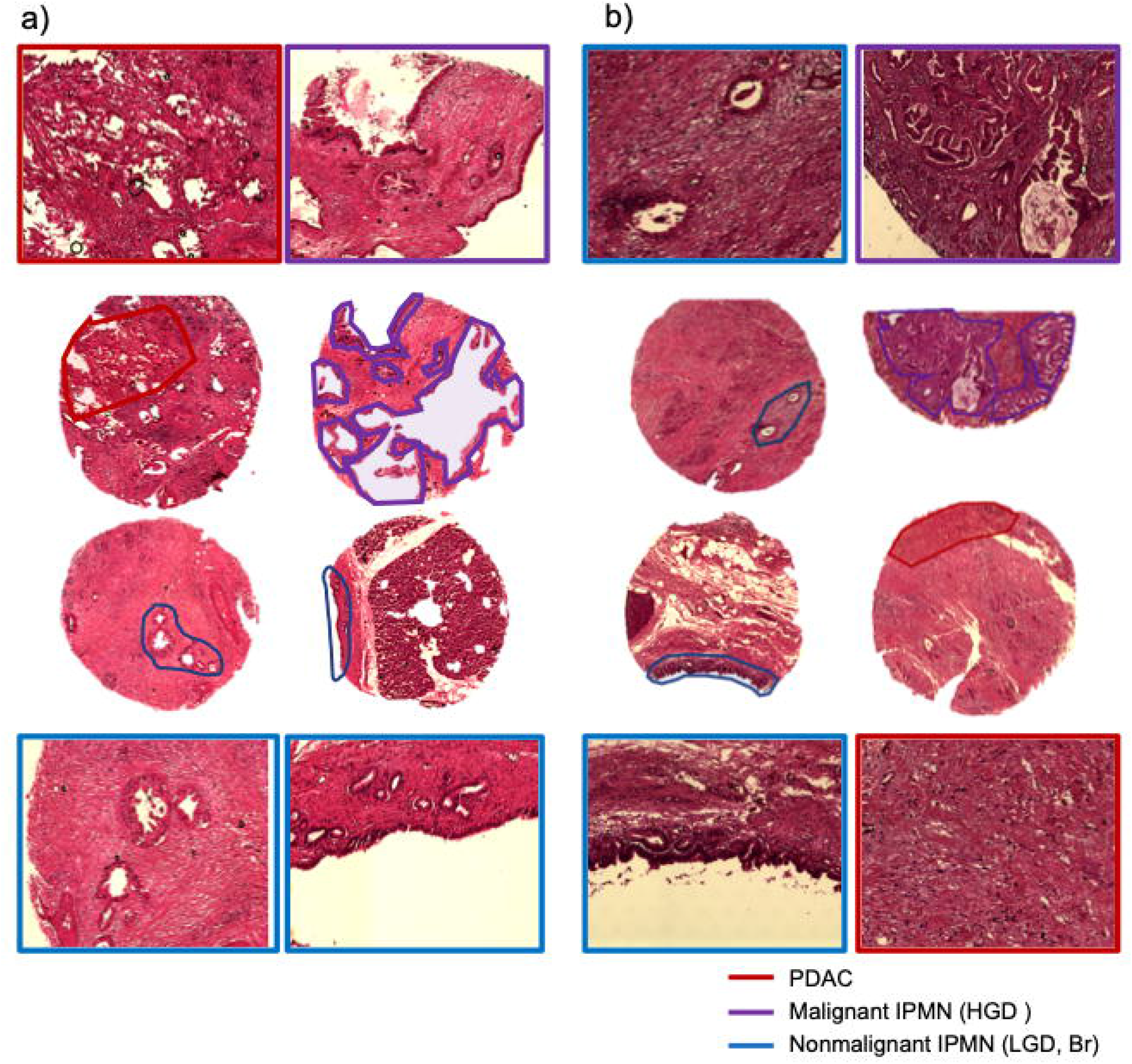
H&E pictures of TMA1 and TMA2. Figure showing TMA 1 and 2 analyzed by Visium Spatial technology. TMA1, LGD (n=2), HGD (n=1) IPMNs and PDAC (n=1); TMA2, LGD (n=1), Br (n=1), HGD (n=1) IPMNs and PDAC (n=1). Red line, PDAC; Purple line, Malignant IPMN (HGD); Blue line, Nonmalignant IPMN (LGD, Br).

An independent validation cohort was obtained by the Australian Pancreatic Cancer Genome Initiative (APCG) consisting of two TMAs (TMA3 and TMA4) with 60 clinically annotated IPMN divided in LGD, borderline, and HGD. This TMA was used for Spatial Transcriptomics with the Nanostring GeoMx Digital Spatial Profiler available in our institution. TMA3 and TMA4 slides were conserved at −80°C enclosed in a sealed bag with silica gel beads prior to shipping in dry ice.

### Spatial Transcriptomics

#### Visium Spatial

TMA1 and TMA2 were used for Visium Spatial Transcriptomics using Visium Spatial for FFPE Gene Expression Starter Kit, Human Transcriptome (10X Genomics,USA) following the manufacturer protocols and recommended third party reagents and supplies. Visium spatial libraries were sequenced with NextSeq 550 (Illumina, USA) at a coverage of 140 million reads in paired-end for each capture area according to manufacturer indications.

BCL files were obtained and processed with the Space Ranger Pipeline from 10X Genomics. Space Ranger output files were imported in R with STUtility (13) package and mapped to the reference H&E stained picture. The imported outputs from each capture area were converted in a Seurat object (14) and integrated using the Harmony algorithm (15). The integrated dataset was analyzed with Seurat and clustering was performed using the Leiden algorithm. Markers from each cluster were identified with FindMarkers() function and for Enrichment Analysis with enrichR interrogating Panglao and Celltype Augmented 2021 datasets to identify cell type belonging to each cluster. These results were integrated with the prediction obtained from the RunAzimuth() function. Differential Expression Analysis (DEA) between IPMN clusters was performed using FindMarkers() function whose output was used for Gene Set Enrichment Analysis (GSEA) with the R package clusteRprofiler interrogating the MsigDB database. To confirm the clusters obtained with Seurat and perform Spatial trajectory inference analysis we used the stLearn (16) Python library. Again Space Ranger outputs were imported in Python and integrated with the Python module for the Harmony algorithm. Subsequently, integrated data was clustered using the leiden algorithm and Spatial trajectory inference analysis was performed.

#### GeoMx Spatial

IPMN TMA 3 and 4 obtained from APCG were analyzed for Spatial Transcriptomics using GeoMx Human Whole Transcriptome Atlas (Nanostring, USA) following the provided protocol. TMAs were stained with GeoMx morphology kit to mark neoplastic cells (PanCK), and stroma cells (CD45). 60 ROI were selected and segmentation was performed to isolate only the PanCK positive area to obtain only the IPMN specific transcriptome. GeoMx library was sequenced with NovaSeq 6000 at a coverage of 541 million of reads. Sequencing data was uploaded on Illumina BaseSpace hub and processed with DRAGEN to obtain .dcc files. The files were imported on R with the GeoMxTools R package and quality control (QC) was performed using default parameters. Only the 43 ROIs (6 LG, 11 Borderline, and 26 HG) that passed the QC were normalized, converted in a Seurat object and used for further analyses and data visualization.

## Results

### Spatial transcriptomics QC validation

IPMN tissues from patients subjected to surgical resection were collected (Fig. 1).

Low-grade and borderline IPMNs were collected from patients that have not shown disease progression for more than ten years and were considered nonmalignant for analyzes.

To our knowledge, this is the best method to take in consideration the transformation time and therefore the real malignancy potential.

Tissues with high-grade dyspalsia associated with PDAC were considered malignant IPMNs. The tissues were reviewed by an experienced pathologist choosing the epithelial structure of the cysts that were considered for the construction of the TMAs.

The cysts included in the TMAs derive from a single patient (totyal = 8) and are considered representative of that patient to take into account inter-patient heterogeneity.

Two TMAs were analyzed by Visium Spatial technology (10X, US) (Fig. 1). TMA1 contained LGD (n=2), HGD (n=1) IPMNs and PDAC (n=1); TMA2 contained LGD (n=1), Br (n=1), HGD (n=1) IPMNs and PDAC (n=1). All tissues were reviewed by an expert pathologist that confirmed the grade and a gastric-like histology for HGD IPMNs.

A total of 1,586 spots were covered for TMA1 with a mean of 19,795 reads per spot and 1,969 genes per spot. Similarly, 1,737 spots were covered for TMA2 with a mean of 19,705 reads per spot and 1,843 genes per spot. These results are good especially if we take into account that most of the spots covered areas with low cellularity and full of connective tissue.

Unfortunately, although visium has cutting-edge resolution (55μm) we lost almost one entire LG-IPMN in TMA1 wich monolayer epithelium ended on the gap between two lines of spots.

To confirm the quality of data and that the tissue preserved architecture during the Visium protocol we assessed the spatial expression of several IPMN and PDAC clinical markers whose expression highlighted only the areas where these were located (Fig. 2a); moreover, low levels of CDX2 and the absence of MUC2 confirmed the gastric-like histology of the IPMNs.

**Figure 2.**
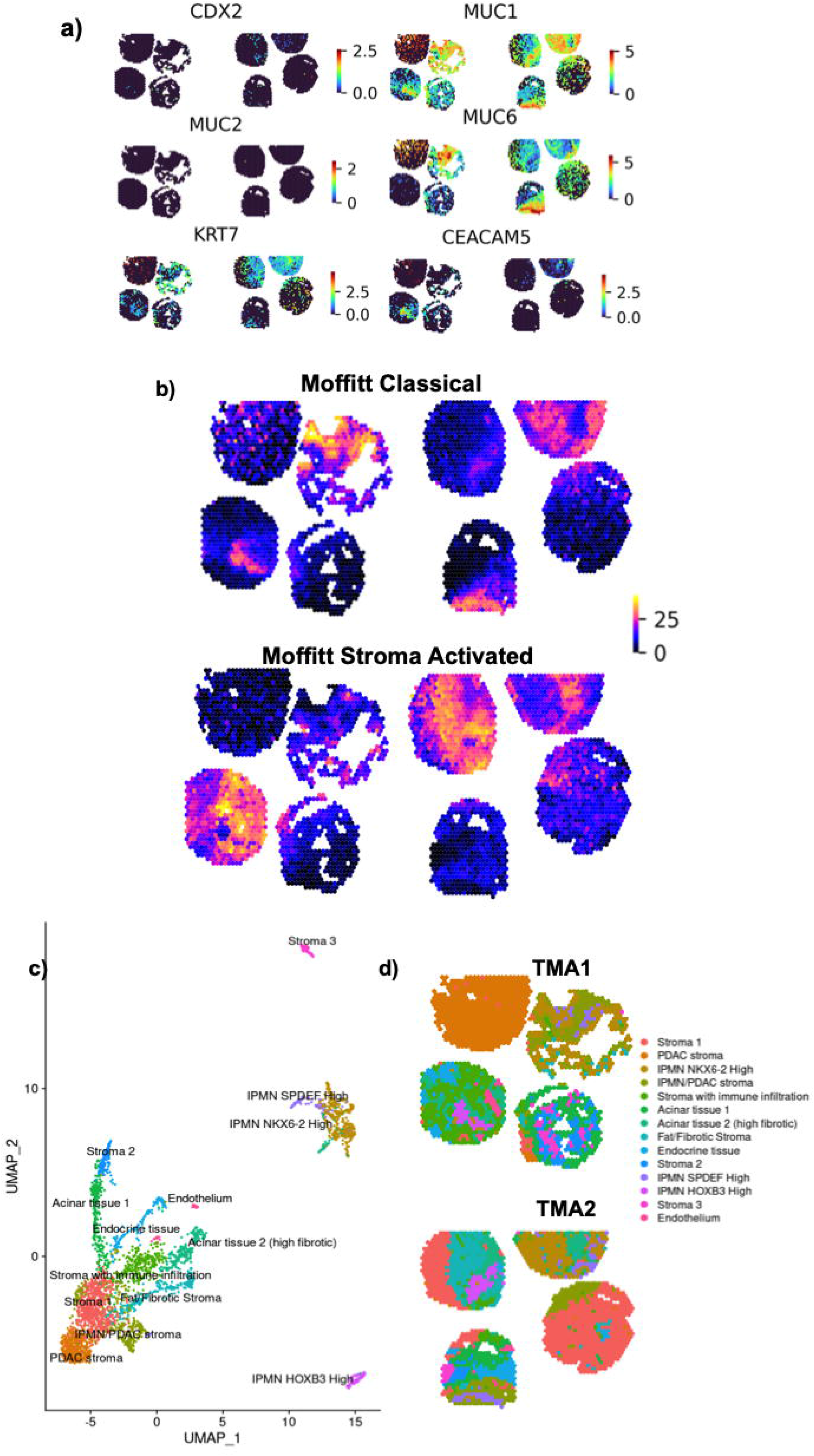
Routine markers, Moffitt Molecular signatures, and Seurat clustering analysis. a) Gene Expression of IPMN routine diagnostic pathology markers by Visium Spatial transcriptomics. Log2 normalized raw counts are shown b) Moffitt gene signatures expression in the TMAs. Cumulative Sum of the gene sets is shown. c) UMAP plot showing the different clusters identified with Seurat and d) their location in a spatial context.

To further confirm the correspondence between the clusters and cell types, we checked the expression levels of the most reliable molecular signatures in PDAC, namely Moffitt signatures (Fig. 2b). As expected, while the tumoral signatures (Classical and Basal) highlighted only the IPMN/PDAC areas, the stromal signatures (Activated and Normal) mapped specifically only in stromal regions.

Interestingly, we found that Classical and Stroma Activated signatures were shared by all samples and were already highly expressed in LGD IPMN (Fig. 2b). We took all these results as proof of the success of Visum’s analysis, so we proceeded with more in-depth investigations.

A total of 14 clusters were identified by bioinformatics analyses (Fig. 2c,d): five different stroma clusters representing different admixtures of mesenchymal cells (Fig. 2c), three clusters representing the pancreatic tissue (acinar and endocrine), three clusters arising in the different types of IPMN, two representing the IPMN and PDAC stroma, and one for endothelium.

Interestingly while LGD IPMNs is a well-separated cluster, Br and HGD IPMNs share a large part of the transcriptome forming very close clusters (Fig. 2c). Visium technology proved to be an effective technology to study small lesions on archived patient tissues. We identified cell clusters that not only mirrored the tissue architecture but that also express common markers of PDAC malignancy.

### Characterization of the three IPMN clusters

To find specific markers of IPMN malignancy we deeply investigated the transcriptome of the three IPMN clusters. Using Seurat, we found the specific markers that characterized the three IPMN clusters and we managed to identify gene signatures that spatially marked the different types of IPMN (Fig. 3a). We found that HGD IPMN (Fig. 3b) were characterized by a high expression of *NKX6-2* (log2FC= 21.9, pvalue= 2.60e-263) the borderline IPMN (Fig. 3c) showed a high and quite specific expression of *SPDEF* (log2FC = 8.6, pvalue= 6.68e-95), and LGD IPMNs (Fig. 3d) had high expression of *HOXB3* (log2FC= 5.1, pvalue= 3.19e-67).

**Figure 3.**
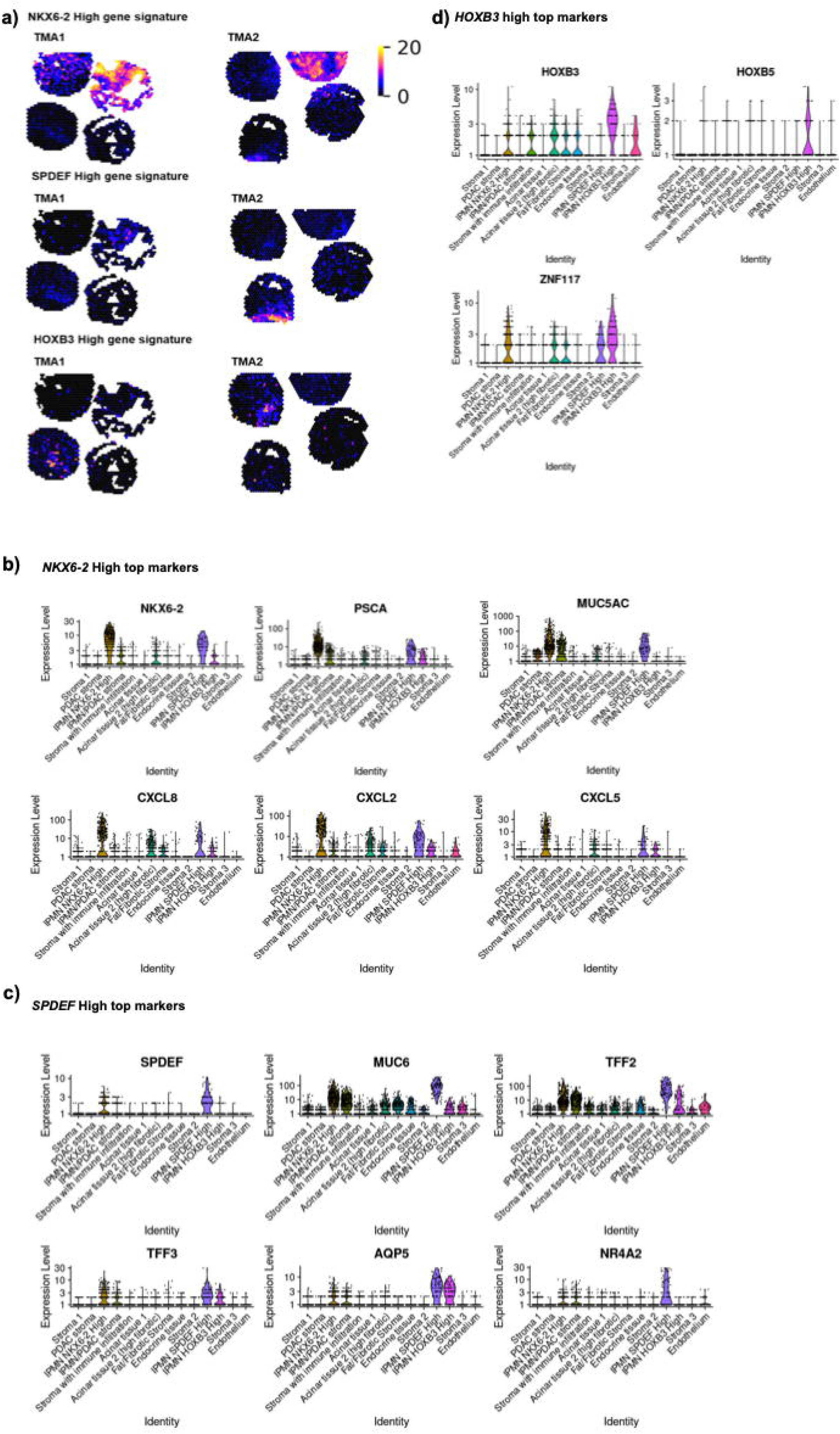
Gene signatures and Markers identified in IPMN clusters. a) Cumulative Sum of the gene signatures identified in *NKX6-2* high, SPDEF high, and *HOXB3* high IPMN clusters. b), c), and d) Violinplot showing the log normalized expression of the top markers for each of the three IPMN clusters.

The NKX6-2 high top markers were genes highly expressed in gastric cells such as *PSCA* and *MUC5AC,* as well cytokines that are associated with cancer progression and immune evasion such as *CXCL2, CXL5, CXCL8* (Fig. 3b).

The SPDEF high IPMN cluster instead was characterized by markers of goblet cells of different tissues such as lung (*SPDEF, TFF2, AQP5*), intestine (TFF3), and gastric (MUC6) (Fig. 3c).

The HOXB3 cluster main markers were 3 transcription factors *HOXB3, HOXB5,* and *ZNF117* (Fig. 3d).

To get more info into the pathogenesis of IPMN we performed DEA and GSEA analysis between *NKX6-2* high and *HOXB3* high IPMN clusters to identify the main pathways activated in the HGD in respect to the LGD IPMN (Fig. 4).

**Figure 4.**
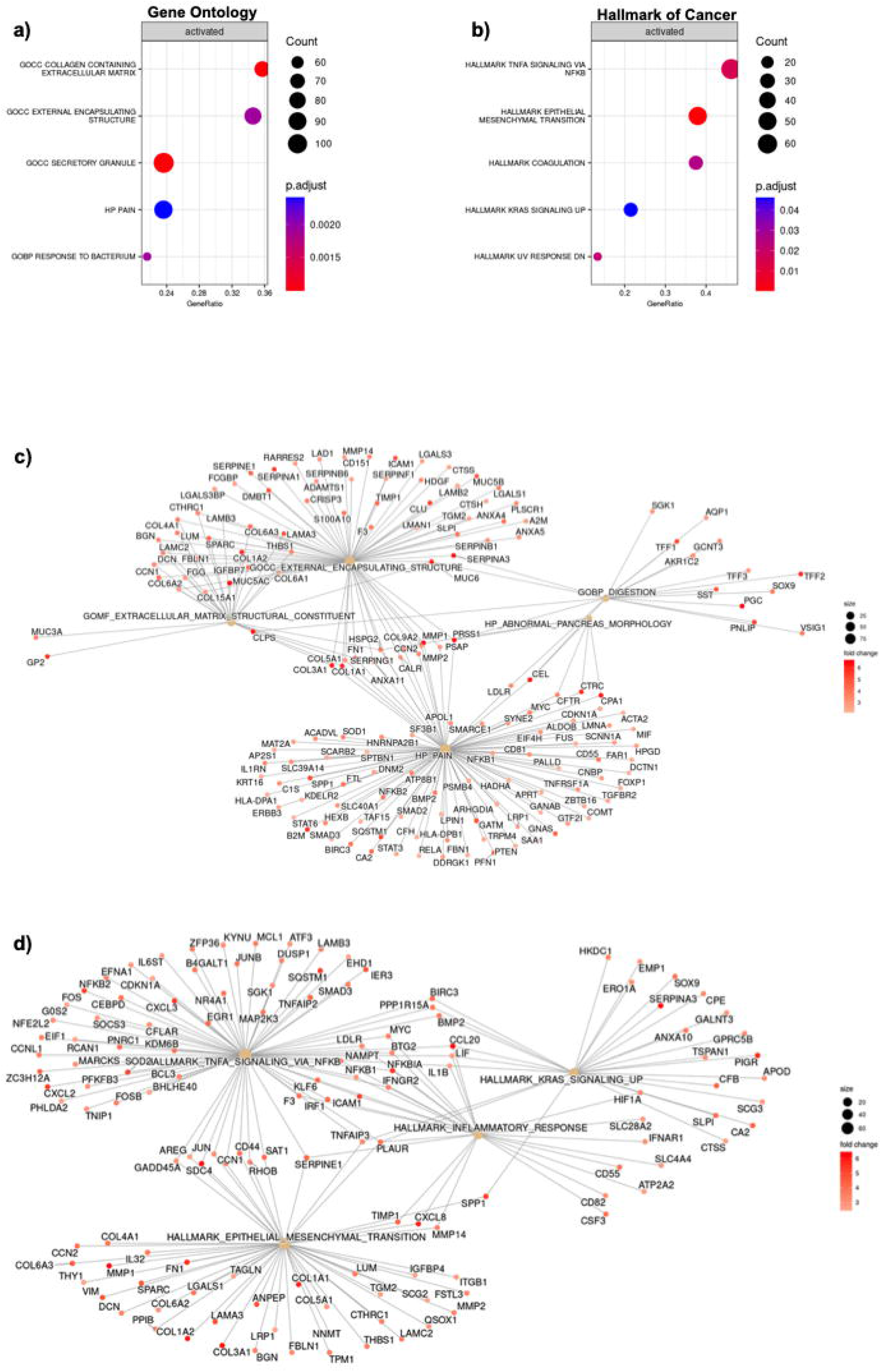
Pathways activated in *NKX6-2* high IPMN in respect to *HOXB3* high IPMN. a) Dotplot describing Gene Ontology GSEA results showing the top 5 activated geneset in *NKX6-2* high IPMN compared to *HOXB3* high IPMN. Dot size indicates the number of genes involved in the geneset, and the color corresponds to the p value. b) Dotplot showing the top 5 activated Cancer Hallmark pathways in *NKX6-2* high IPMN. c) and d) Cnetplot showing the hub genes in the activated genesets. Fold change values were scaled to improve visualization.

Top two Gene Ontology gene sets (Fig. 4a) identified to be upregulated in HGD IPMN involved the extracellular matrix deposition and organization including a compendium of ECM genes actively involved in PDAC oncogenesis such as *FN1, COL1A1, SPARC, MUC5AC, MUC5B, LGALS1, LGALS3, DSC* and many other (Fig. 4c). Another interesting signature HP_PAIN (Fig. 4c) that includes several genes involved in response to tissue damage hinting, together with the other signatures, that *NKX6-2* high IPMN profoundly remodel and affect the pancreatic tissue.

We also interrogated the MsigDB Cancer Hallmark signatures compendium and found that *NKX6-2* IPMN display a consistent activation of *KRAS* and *NFKB* signaling, Epithelial to Mesenchymal Transition, and and a consistent upregulation of genes involved in the regulation of inflammation and immune response such as *CCL20, CXCL2, CXCL5, CXCL8, IL1B,* and *ICAM1.*

Moreover, using the MsigDB Cell Type database we found that *NKX6-2* high IPMN overexpressed genes such as *NKX6-2, S100A14, TFF1, FER1L6* that were identified by Busslinger et al. (17) as the top markers of gastric isthmus cell performing single-cell analyses (Suppl.Fig. 1a)

All these signatures indicate that *NKX6-2* high IPMN activates several and important hallmarks in cancerogenesis and that the gastric signature associated may be a good marker of IPMN malignancy. We also performed DEA between *NKX6-2* high IPMN and *SPDEF* high IPMN and found a total of 177 differentially expressed genes. GSEA analysis showed that most of the genes upregulated in *NKX6-2* high IPMN were involved in the same pathway resulted from the comparison with *HOXB3* high IPMN such as epithelial to mesenchymal transition, extracellular matrix organization, and external encapsulation organization (Sup. Fig. 1b, c).

These findings shed more light into IPMN progression identifying the key genes and pathways that drive the IPMN degeneration to PDAC. GSEA showed that *NKX6-2* high IPMN possesses most of the hallmark signatures that characterize PDAC; KRAS pathway upregulation in particular. Also, the consistent upregulation of genes involved in cell extrinsic pathways (e.g. Inflammation, External Encapsulation, ECM deposition) entails that *NKX6-2* high IPMN intently shape the tumor microenvironment to support its growth activating the same core mechanisms that PDAC exploits for its sustainment.

### *SPDEF* high IPMN evolution in *NKX6-2* high IPMN

To further validate the clusters identified with Seurat we analyzed Visium data also with the stLearn Python package that also implemented an algorithm for spatial trajectory inference. The stLearn package identified a similar cluster to Seurat and also confirmed the three IPMN clusters (Fig. 5a, b).

**Figure 5.**
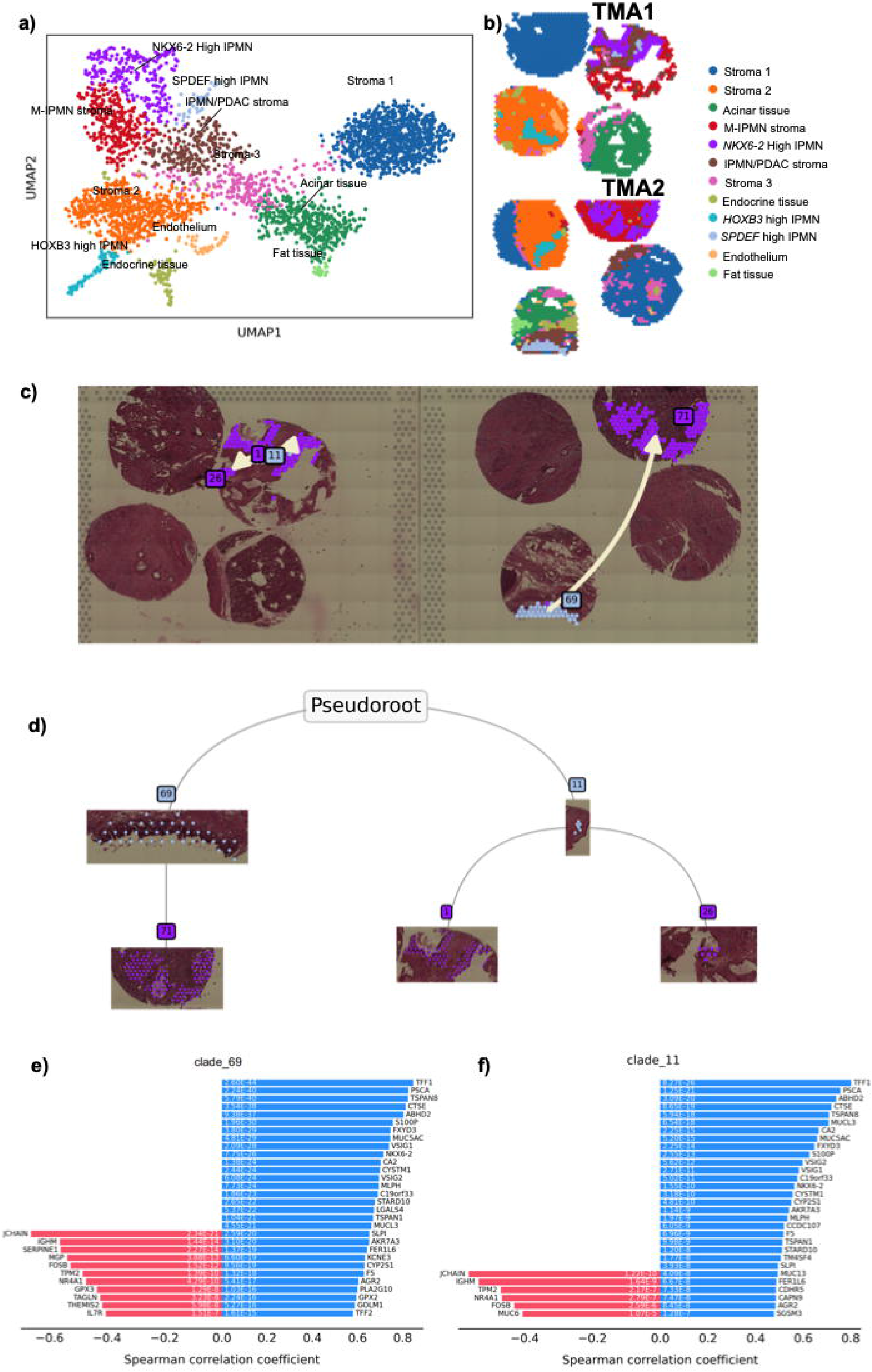
Spatial trajectory analysis. a) UMAP plot showing the cluster identified with stLearn and b) their location in the analyzed tissues. c) and d) Spatial trajectory tracing the evolution from *SPDEF* high to *NKX6-2* high IPMN in both the TMAs analyzed. e) and f) Barplot showing the genes that correlate positively (blue) and negatively (red) with the spatial trajectory.

Since the *SPDEF* high and *NKX6-2* high IPMN clusters were similar at transcriptomic level (Fig. 2a, Fig. 5a). We performed the spatial trajectory analysis to investigate the possibility that *SPDEF* high IPMN cluster represents a Br IPMN that is likely to evolve into a malignant HGD *NKX6-2* high IPMN.

The analysis actually confirmed this hypothesis and showed how *NKX6-2* may drive this evolution as it is the only transcription factor included in the gene list of genes that correlates with the trajectory. Most of the other genes in the top tierlist such as *PSCA, TFF1, TFF2, MUC5AC, MUCL3, FER1L6* are gene that are typical markers of gastric cells hinting that the acquisition of a gastric-like morphology is a marker of an acquired malignant potential.

### Validation of the identified markers of malignancy

To understand if the markers identified with Visium maintained their significance in a more macroscopic context we used another spatial approach using Nanostring GeoMx technology on an independent series of 43 IPMN consisting of 6 LGD IPMN, 11 Br IPMN, and 26 HGD IPMN. We stained TMAs using GeoMx morphology kit to mark neoplastic cells (PanCK), and stroma cells (CD45), to perform cell segmentation to isolate on the IPMN transcriptome (Panck+) and discard the stromal area (CD45+) (Fig. 6a). We selected ROIs including 500-800 cells or, as in the case of LGD and BrIPMNs, the whole lesions (Fig. 4b). These analyses validated the Visium results where most of the markers identified such as *NKX6-2, SPDEF, MUC5AC,* and *PSCA* were found upregulated in TMA3 and TMA3 HGD IPMNs. We also assessed the gene set activity of the *NKX6-2 high* signature in the validation TMA cohort using the Seurat function AddModuleScore() Fig nanostring and we found that HGD IPMN had the highest score while LGD IPMN showed negative scores indicating the absence of expression of such genes.

**Figure 6.**
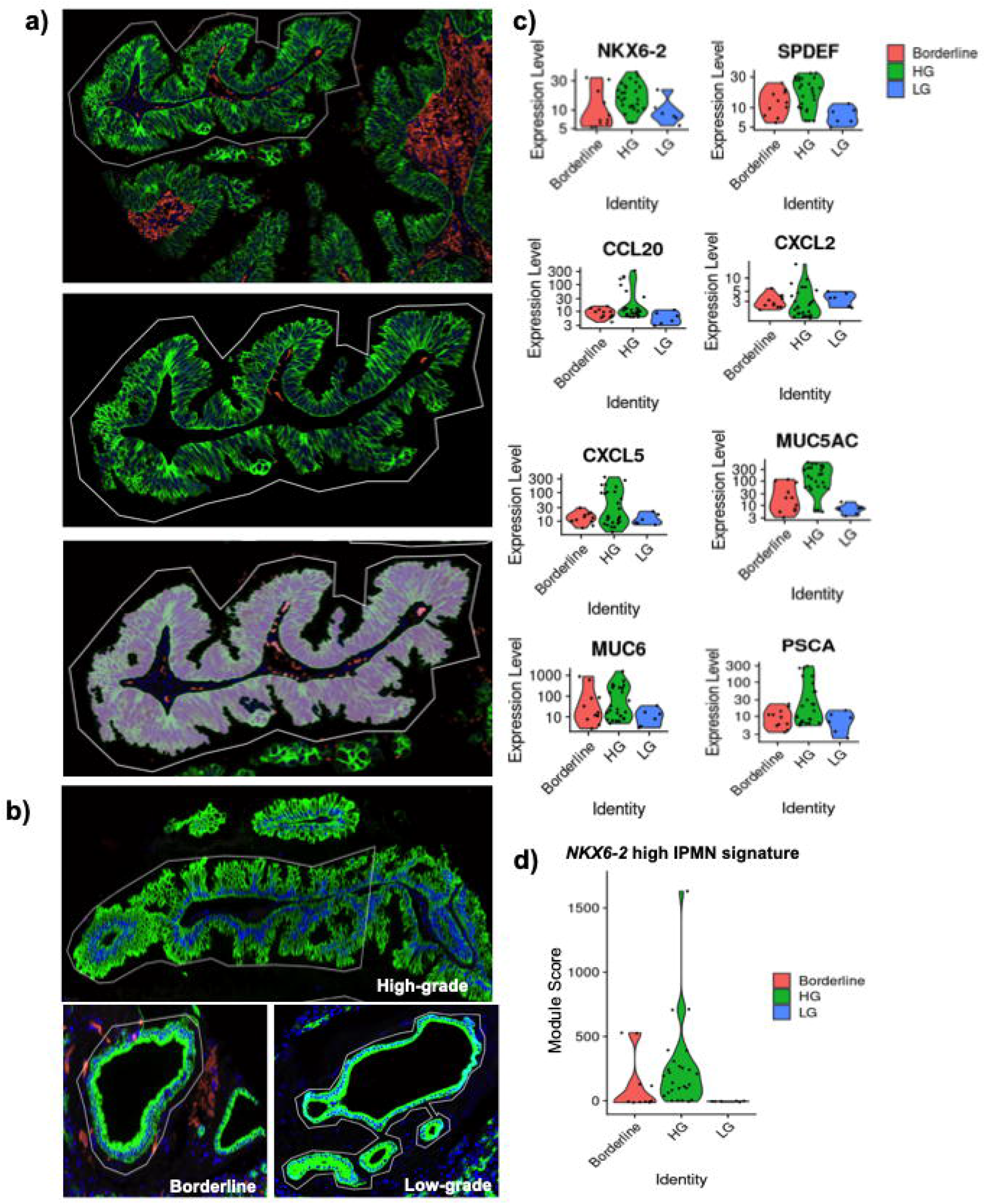
GeoMx spatial transcriptomics analysis. a) Pictures showing ROI selection and segmentation strategy chosen for the analysis. b) Picture showing three ROIs representative of the histology group selected LGD, Br, and HGD. c) Top markers identified in HGD IPMNs by GeoMx analysis. Log transformed normalized counts are shown. d) *NKX6-2* high IPMN gene set activity calculated with Seurat AddModuleScore() function for the three IPMNs groups.

With this analysis we were able to validate the malignancy markers associated with Gastric phenotype that we have found with Visium, strengthening the evidence that the gastric transcription program is a hallmark of IPMN degeneration toward PDAC.

## Discussion

Intraductal papillary mucinous neoplasm (IPMN) are mucin-producing cysts classified as premalignant lesions. The high-risk IPMNs are addressed to surgery (57-90%), while low-risk IPMNs (6-46%) are referred to imaging for monitoring of the development of malignant features. Unfortunately, a little part of the cysts (1-11%) that do not undergo surgery progresses into invasive carcinoma (18).

Therefore, the identification of the molecular events driving the progression of premalignant lesions in full-fledged carcinoma would serve the identification of interception and prevention strategies. In this study we used two spatially resolved transcriptomics to characterize the epithelial compartment of malignant and nonmalignant IPMNs in order to uncover new biomarkers for risk stratification of IPMN patients as well as mechanisms potentially responsible for malignant transformation.

This work is based on the new technological advances of Next-Generation-Sequencing combined with imaging that have reached a very high resolving power allowing to measure the levels of expression of all genes systematically and to spatially resolve and associate gene expression to the specific cell (or group of cells) or tissue architecture.

Iyer and colleagues, using spatial transcriptomics analysis (GeoMx, Nanostring) identified a molecular framework for risk stratification of IPMN patients (19). In this interesting preprint paper, only the 10% of transcriptomes of IPMNs samples (n=12) were analyzed, each containing both different areas of LGD and HGD IPMNs.

Thus, it is conceivable that the LGD from samples completely devoid of HGD and malignant tissues may differ completely at transcriptomics level.

Numerous papers proposed mutational aberration (20), extravesical (EV) proteins (21) and proteins (22), as the main transforming factors of IPMN.

However all the attempts to translate into clinical settings the putative identified markers of IPMN malignancy were disappointing and it is still an urgent clinical need.

Mainly this is due to the complexity of the pathology, with very small heterogeneous cysts and to the limited technologies both for the number of analytes and for the type of analyzed tissues. All these papers based the analysis exclusively on IPMN dysplasia grade or IPMN diverse epithelial subtypes without taking into account the true potential for malignant transformation of IPMNs (23–27).

Our work is aimed at identifying markers for the clinical stratification of IPMN patients and is based not only on the different IPMNs grade but also on the temporal criterion that includes patients with clinical follow-up greater than 10 years who did not experienced disease recurrence.

The gene signatures identified in this study can distinguish between Non-Malignant and all Malignant IPMNs pinpointing on three transcriptional factors specifically expressed in each IPMN stage: *HOXB3* (LGD, Nonmalignant), *SPDEF* (Br Nonmalignant) and *NKX6-2* (HGD Malignant).

SPDEF is an established regulator of secretory cell differentiation during development (28,29). In particular, Tonelli C., using a PDAC progression mouse model identified Spdef in an epithelial-high cell subpopulation, by scRNA-seq analysis, as a factor required for tumorigenesis in pancreatic cancer cells of epithelial and mucinous nature (30). This exciting paper reinforces our evidence, considering that the model uses KRAS and P53del expression throughout the pancreas (PDX1), it is conceivable that many more mucinous neoplastic epithelial cells were included in the early stages of carcinogenesis.

The exact carcinogenesis of PDAC is still under investigation, and the generic notion that the duct gives rise to IPMN while the acinar cell to PanIN is weak and discussed. This is highlighted by several studies in which the transition from acinar to duct cells is well demonstrated, as well as the existence of intimal interconnection between differentiated cells capable of dedifferentiate acquiring the properties of more immature cells within the same lineage hierarchy (31).

*NKX6-2* is a transcription factor acting on pancreas embriogenesis and specifically on endocrine progenitor cell differentiation (32). Its expression is common to cells of gastric isthmus (Suppl.Fig 1a) and gastric subtype IPMNs (17) and reinforcing the hypothesis of “paligenosis” from gastric and pancreatic ductal tissues (33).

*HOXB3* expression has been associated with normal characteristics and many papers show that *HOXB3* silencing leads to a less aggressive phenotype of tumor cells by restoring epithelial characteristics of tumor cells (34–36). It has also been reported a simultaneous hypermethylation of HOX genes, including HOXB3, in pancreatic cancer. Here we showed that LGD Non Malignant IPMN has a HOXB3 signature with respect to IPMN with malignancy features.

Cell trajectory analysis revealed that IPMN components activate multiple transcription factors differentially along histology progression that seems to delineate a continuum between Br - and HGD Malignant-IPMNs.

The marker identified by Visium were confirmed, using GeoMx Digital Spatial Profiler, in an external validation cohort (n = 43) provided by the APGI and consisting of patients with different IPMN histologies. Even Though, the spatial resolution of this technology is lower than Visium, it is still a valid and robust method to precisely characterize the transcriptome of small regions of tissue. As expected, the analyses on GeoMx spatial transcriptomics data confirmed the association between the *NKX6-2* high IPMN signatures and tumor progression.

## Conclusions

Overall, our results provide valuable resources for deciphering the gene expression landscapes of

IPMN cysts. Meanwhile, we revealed, and validated in an independent cohort of IPMN patients, critical signaling pathways and transcription factors that could coordinate the transition from LGD non-malignant to IPMN with malignant features.

Therefore, these findings are potentially useful in advancing our current understanding of the critical genetic network related to PDAC progression, laying the foundation for a preventive analysis of IPMNs and for initiating patients to various therapeutic interventions.

## Acknowledgements

We thank the Australian Pancreatic Cancer Initiative (APGI) for sharing IPMN derived patient tissues; 10X Clinical Translational Research Network (CTRN); Fondazione Nadia Valsecchi Onlus and Italian pancreatic cancer Community (IPCC).

## Ethics declarations

The experimental protocol was approved by the local ethics committee (Fondazione Policlinico Gemelli IRCCs, Ethical Committee approval Prot. Gen. 3536) and followed EU regulations.

## Consent for publication

Not applicable.

---

**Supplementary Figure 1.**
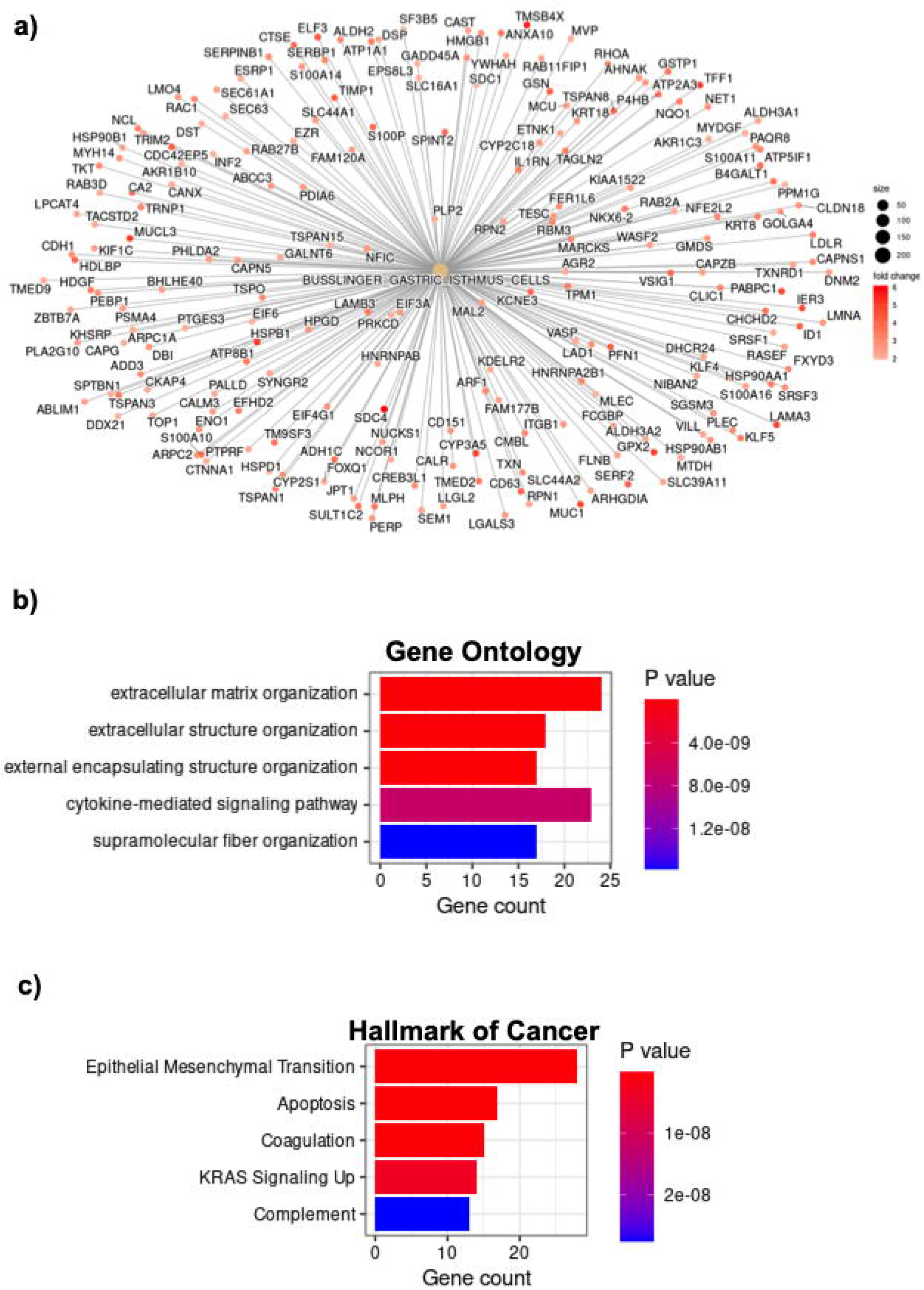
GSEA analysis additional figures. a) Cnetplot showing the gastric Isthmus cell signature that was found upregulated in *NKX6-2* high IPMN compared to *HOXB3* high IPMN. Fold change values were scaled to improve visualization. Barplot showing the Gene Ontology b) and Hallmark of Cancer c) gene sets overexpressed in *NKX6-2* high IPMN in respect to *SPDEF* high IPMN.

